# At a snail’s pace: the influence of habitat disturbance on terrestrial snail movement using experimentally manipulated mesocosms

**DOI:** 10.1101/2021.10.05.463224

**Authors:** Emily E. Denief, Julie W. Turner, Christina M. Prokopenko, Alec L. Robitaille, Eric Vander Wal

## Abstract

The Anthropocene marks great changes to environments and the animals that inhabit them. Changes, such as disturbance, can affect the manner in which animals interact with their environments, such as moving and selecting habitats. To test how animals might respond to changing disturbance regimes, we employ an experimental approach to movement ecology. We used integrated step selection analysis (iSSA) to test the behavioural responses of individually-marked grove snails (*Cepaea nemoralis*) exposed to a gradient of physical disturbance in their habitat. We used a before-after control-impact (BACI) experimental design within semi-controlled mesocosms to manipulate edge and disturbance variables by altering the area of the mesocosm covered by bricks. We showed that grove snails perceive edges of enclosures and edges of bricks as risks, and responded to such risks by altering their movement. Grove snails displayed a bimodal response to risk by taking shelter in place or moving faster to be farther from the disturbance. Furthermore, individuals tended to modulate their behavioural response to the degree of risk. Our study highlights the usefulness of experimental mesocosms in movement ecology and in determining the effects of habitat alteration and human-imposed risk on movement behaviour.

## Introduction

Threats of climate change and anthropogenic habitat alteration are becoming increasingly prevalent (Nicolai & Ansart, 2017). 50-70% of Earth’s surface has been altered by humans, resulting in changes to habitat composition, biodiversity, and the movement dynamics of many animals (Tucker et al., 2018). The alteration of ecosystems by humans can result in habitat fragmentation, which leads to an increase in the amount of edge habitat in a system. One of the most prominent causes of habitat fragmentation is the construction of roads through habitat patches (Belmeziti et al., 2018). Roads can have many negative effects on the abundance of animals, including disturbances from noise and pollution causing avoidance of the area, and road mortality (Fahrig & Rytwinski, 2009). Roads fragment the habitat creating a larger edge to area ratio (Golden & Crist, 2000). With an increased amount of edge, habitats experience a change in species abundance and composition (Golden & Crist, 2000). Species richness in ants has been shown to be smaller in slightly fragmented habitats as compared to unfragmented habitats, but greater in habitats that have been highly fragmented (Golden & Crist, 2000). Other consequences of habitat alteration include reduced resources and modified dispersal behaviour in a number of species (Frid & Dill, 2002; Nicolai & Ansart, 2017).

Movement is an integral component of many ecological systems. An animal’s movement is associated with various processes and patterns, and thus it should be explicitly considered when conducting studies at the individual, population, and ecosystem level (Nathan, 2008). Changes in movement behaviour can be seen as a response to risk (Frid & Dill, 2002). The risk-disturbance hypothesis (Frid & Dill, 2002) suggests that individuals perceive human disturbance at the same level that they would perceive predation risk. When human disturbance is present in a habitat for an extended period or is intensified, individuals will tend to shift habitats to avoid such disturbance even if this reduces their access to resources (Frid & Dill, 2002). This behaviour can be seen in elk, which respond to roads the same way they would predation risk by selecting for areas farther from roads and seeking cover when near roads (Prokopenko et al., 2017). Similarly, bottlenose dolphins have been recorded avoiding high-quality forage areas when motorboat density in the area was high (Allen & Read, 2000). The theory also predicts that the probability of fleeing to escape risk increases when the costs of fleeing are lower (Frid & Dill, 2002). This cost depends on the quality and distribution of resources, environmental conditions, and the mobility of the animal (Denny, 1980; Frid & Dill, 2002). Animals are predicted to respond more strongly when the disturbance stimuli are larger in size, though this prediction was previously unable to be evaluated (Frid & Dill, 2002).

Gastropods are expected to be very sensitive to habitat alteration (Nicolai & Ansart, 2017). Increased habitat fragmentation has led to modified dispersal behaviours, which have been expressed differently across species of snail. In the rock-dwelling land snail *Albinaria caerulea*, avoidance of fragment edges has been observed (Giokas & Mylonas, 2004). Contrary to this, the brown garden snail *Cornu aspersum* was recorded exhibiting higher levels of movement as fragmentation increased (Dahirel et al., 2016). As predicted by the risk-disturbance hypothesis (Frid & Dill, 2002), we should expect gastropods to respond to human disturbance as they would predation risk. Snails have been recorded exhibiting antipredator behaviour in the presence of both visual and chemical alarm cues (Dalesman et al., 2006). As animals with low mobility and a high cost of movement, it is often ineffective for gastropods to actively escape risky environments (Denny, 1980). Trade-offs between energy cost and predation risk have been observed in the great pond snail *Lymnaea stagnalis*, which only exhibited escape behaviour when risk levels were moderate or high (Dalesman et al., 2006). In situations where escape is not possible or has too high a cost individuals may respond to risk by seeking refuge in habitats that provide cover or, in the case of many snails, retreating into their shell (Hebblewhite & Merrill, 2009; Trussell & Nicklin, 2002). Other responses such as burying themselves in substrate were recorded in aquatic snails that were presented with alarm cues (Aizaki & Yusa, 2009). Many of these studies observed behavioural changes in gastropods when presented with the risk of predation. Though gastropods are more sensitive to human disturbance than many other taxa, there are few studies on the impact it has on them (Nicolai & Ansart, 2017). Integrated step selection analysis (iSSA; Avgar et al., 2016) can be a valuable tool in predicting the effect of human disturbance on selection and movement in animals. Using snails to model this behaviour is valuable, as with a small, low-mobility species it is possible to have a smaller and more manageable study area where factors such as edge density and habitat type can be easily controlled and manipulated experimentally.

Here, we present grove snails (*Cepaea nemoralis*) with varying degrees of risk in the form of edges and physical human disturbance in a controlled experimental mesocosm, and model these predictions using iSSA. We hypothesize that snails are able to perceive edges as risks and respond to these risks by modifying their behaviour. We predict that snails will avoid areas near edges as a way to avoid risk. They will seek refuge by hiding in their shells and reducing movement rates during times of risk when they are near edges. As well, we hypothesize that snails are able to assess the degree of risk and modify their behavioural response as the degree of risk varies. We predict that snails will decrease their movement rates as the area covered by the physical human disturbance within the mesocosm increases.

## Methods

### Study Area

The study took place on 4.5 m^2^ of grass lawn in downtown St. John’s, Newfoundland and Labrador. Data collection occurred from September 26 to November 2, 2019. The temperature during this time averaged ten degrees Celsius. We constructed eight closed mesocosms measuring 65 × 65 × 30 cm using 1 cm wire mesh hardware cloth and wooden frames, which were placed in a two by four array across the study space with 15 cm between each mesocosm. The grass density was estimated to be nearly constant at 21 blades per centimetre squared across the entire area. This was calculated by sampling the number of blades within a 10 × 10 cm quadrat in each mesocosm. The ground was level across all of the mesocosms.

### Study species

The experiment was conducted using grove snails, *Cepaea nemoralis*, a terrestrial snail in the family Helicidae. This species was introduced to North America from Western Europe (Hoxha et al., 2019) and is now abundant in urban areas of Newfoundland. They are nocturnal (Richardson, 1975) and mainly forage on dead or dying plant material (Wolda et al., 1971). The snails are active until late fall, then hibernate over winter buried under leaf litter or soil (Wolda et al., 1971). Snails were collected from areas surrounding downtown St. John’s and housed indoors in a glass terrarium covered with burlap to allow for oxygen flow until moved to the mesocosms for the experiment. While indoors, snails were kept at room temperature and away from direct sunlight. They were provided with water, and foliage from several plants as food. Individuals weighed between 1-3 g. Shells were approximately 2 cm in diameter. Effort was made to use similar-sized individuals, though there was variation in shell colour polymorphisms. New snails were used in each serial replication of the experiment.

### Experimental setup

Each of the eight mesocosms were monitored using a RECONYX™ PC900 HyperFire™ trail camera. Images were taken every 30 minutes for a total of nine days. Two snails were placed within each enclosure and allowed to acclimate for 24 hours while being monitored by the trail cameras. Snails within an enclosure were differentiated by painting their shells different colours and patterns with nail polish. The painting of their shells allowed for the snails to be distinguished in photos both during the day and at night with infrared imaging. A ruler was placed along the edge of each mesocosm so a scale could be set during image processing.

The experiment was run as a Before-After-Control-Impact (BACI) design. This design has been shown to be a powerful tool in assessing the effects of environmental impact and human disturbance on ecosystems (Conner et al., 2016). The “Before” stage started immediately following the 24-hour acclimation stage and lasted a total of 96 hours. During this stage, the ground of the mesocosm was completely covered by grass. The “After” stage introduced three different treatments to the mesocosms to simulate a physical human disturbance, as well as a control. Each of the treatments took place in two mesocosms. The first treatment was adding one brick with an area measuring 10 × 20 cm to the upper right corner of the enclosure, with a total covered area of 200 cm^2^. The second treatment was adding two bricks in the upper right corner, for a total covered area of 400 cm^2^. For the third treatment four bricks were added to the upper right corner, covering a total area of 800 cm^2^. For the control, no bricks were added. Snails were subsequently monitored for 96 hours. Three iterations of the experiment were run serially, each using new snails. Following each iteration images were downloaded from the trail cameras. Images were analyzed using ImageJ software (version 2.0.0-rc-69/1.52p) to determine the coordinates of each snail on a cartesian grid. We obtained temperature data from Environment and Climate Change Canada.

### Integrated step selection analysis (iSSA)

Resource selection analysis (RSA) offers an intuitive approach to estimate the probability of a spatial resource unit being used based on the type and value of the resource (Manly et al., 2002). Though effective in explaining spatial population distribution based on resources, RSAs often do not consider the accessibility of spatial units (Matthiopoulos, 2003). Step selection analysis (SSA) is a movement-informed RSA that addresses this problem by measuring used steps against available steps. In these models, steps are defined as a straight line connecting consecutive locations of an individual and available steps are sampled from an empirical distribution of used steps based on their length and turn angle (Fortin et al., 2005). A relatively new method of exploring movement and selection is integrated step selection analysis (iSSA; (Avgar et al., 2016). iSSA allows for stronger predictions of space use through the simultaneous estimation of both movement and selection parameters (Avgar et al., 2016). Here, we followed the method for individual-specific RSAs described by Muff et al. (Muff et al., 2019), running the model as a Poisson point process regression with the package *glmmTMB* (Brooks et al., 2017).

All analyses were conducted using R version 4.1.0 (R Core Team, 2021). To create ten available steps for each used step, step lengths and turn angles were randomly sampled from the distribution of used steps. Within the model step length is ln-transformed, resulting in the covariate being a modifier of the shape parameter of the gamma distribution (Avgar et al., 2016). The cosine of the turn angle was used to transform the covariate into a linear measurement, where positive values denote forward movement and negative values denote backwards movement. Snail tracks, their random steps, and covariates were extracted using the *amt* package (Signer et al., 2019). For the control enclosures, a “ghost brick” proximity raster was added as another control to test if snails are reacting to the corner even when a brick has not been placed there. Snails missing data for more than 20 steps were excluded from the analysis. The final sample size was 24 (control = 6, 1 brick = 6, 2 bricks = 7, and 4 bricks = 5) individuals.

The model included core covariates to control for the ecology of the study species: time of day and temperature. Grove snails are nocturnal (Richardson, 1975) and were expected to move more and forage during the night. We obtained sunrise and sunset times for the study period from National Research Council Canada. “Night” was classified as the hours between sunset and sunrise, and “Day” as the hours between sunrise and sunset. Terrestrial gastropods may enter a dormant state and decrease activity below freezing temperatures (Nicolai & Ansart, 2017), hence the inclusion of temperature.

In addition to the core covariates, we included disturbance covariates to test our hypotheses. In the first model, we included distance from both the edge of the brick and the edges of the enclosure at the end of the step to test the prediction that snails will avoid risk by selecting for areas farther from edges. Furthermore, we predicted that movement speed would be reduced as the level of risk increases; thus, in our second model, we also distance from the edge of the brick and enclosure at the beginning of each step with the natural log of step length to ascertain how the distance to edge affects movement. To account for the BACI design, we interacted all covariates with the treatment (1, 2, or 4 bricks or ghost bricks) by stage (“Before” or “After”). Each covariate had an associated random effect in the regression to control for individual differences.

### Binomial movement analysis

While collecting the data, we observed that some individuals stopped moving during the experiment. iSSAs require movement to determine space-use, which is why we excluded individuals that had less than 20 steps. However, to fully test how snails respond to disturbance, we added a binomial regression testing if snails moved or not post-disturbance. This model included treatment (1, 2, or 4 bricks or ghost bricks), stage (“Before” or “After”), and their interaction as well as temperature. We included a random effect of ID to control for multiple observations.

## Results

There were 18 snails that were visible and did not escape prior to when they experienced “Before” and “After” stages (n = 6 undisturbed, n =12 disturbed). Snails moved significantly more when temperatures were warmer (β = 0.072, z = 4.91, p < 0.0001, Table 1). When controlling for temperature, snails were 8% more likely to move “Before” than “After” for all treatments (β = -1.25, z = -5.21, p < 0.0001). However, those snails that were exposed to one brick were approximately 8% more likely to move “After” than control individuals (β = 1.63, z = 4.02, p < 0.0001, Fig. 1). Snails that were exposed to two bricks did not significantly differ from control, although they did move less than “Before (β = -0.13, z = -0.32, p = 0.75, Fig. 1). Finally, snails exposed to four bricks were approximately 5% less likely to move “After” compared to control individuals and significantly less than themselves “Before” (β = -1.18, z = -2.68, p = 0.0074, Fig. 1).

**Table 1.**
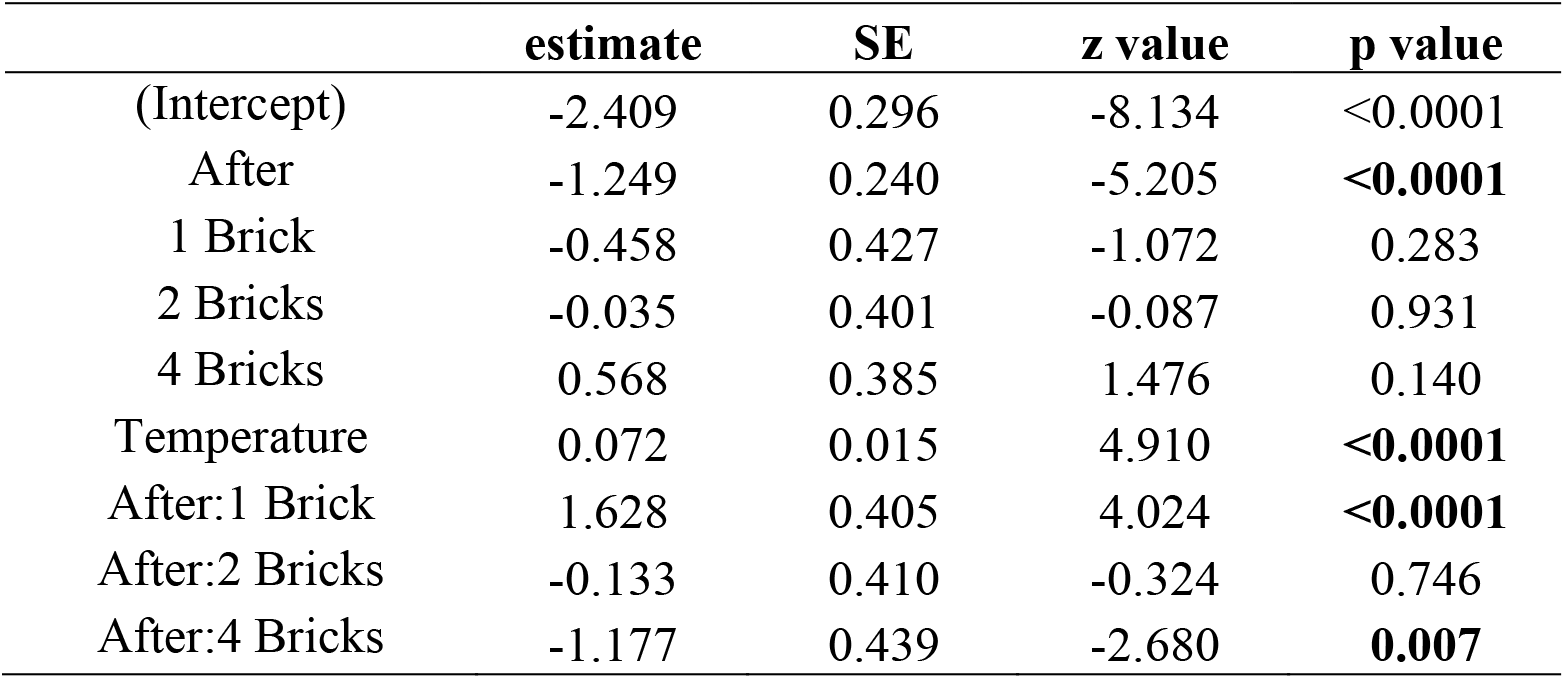
Output from a binomial regression testing the probability of moving or not “Before” and “After” the disturbance treatment. Bolded values are significant at p < 0.05

|  | estimate | SE | z value | p value |
| --- | --- | --- | --- | --- |
| (Intercept) | -2.409 | 0.296 | -8.134 | <0.0001 |
| After | -1.249 | 0.240 | -5.205 | **<0.0001** |
| 1 Brick | -0.458 | 0.427 | -1.072 | 0.283 |
| 2 Bricks | -0.035 | 0.401 | -0.087 | 0.931 |
| 4 Bricks | 0.568 | 0.385 | 1.476 | 0.140 |
| Temperature | 0.072 | 0.015 | 4.910 | **<0.0001** |
| After:1 Brick | 1.628 | 0.405 | 4.024 | **<0.0001** |
| After:2 Bricks | -0.133 | 0.410 | -0.324 | 0.746 |
| After:4 Bricks | -1.177 | 0.439 | -2.680 | **0.007** |

**Figure 1.**
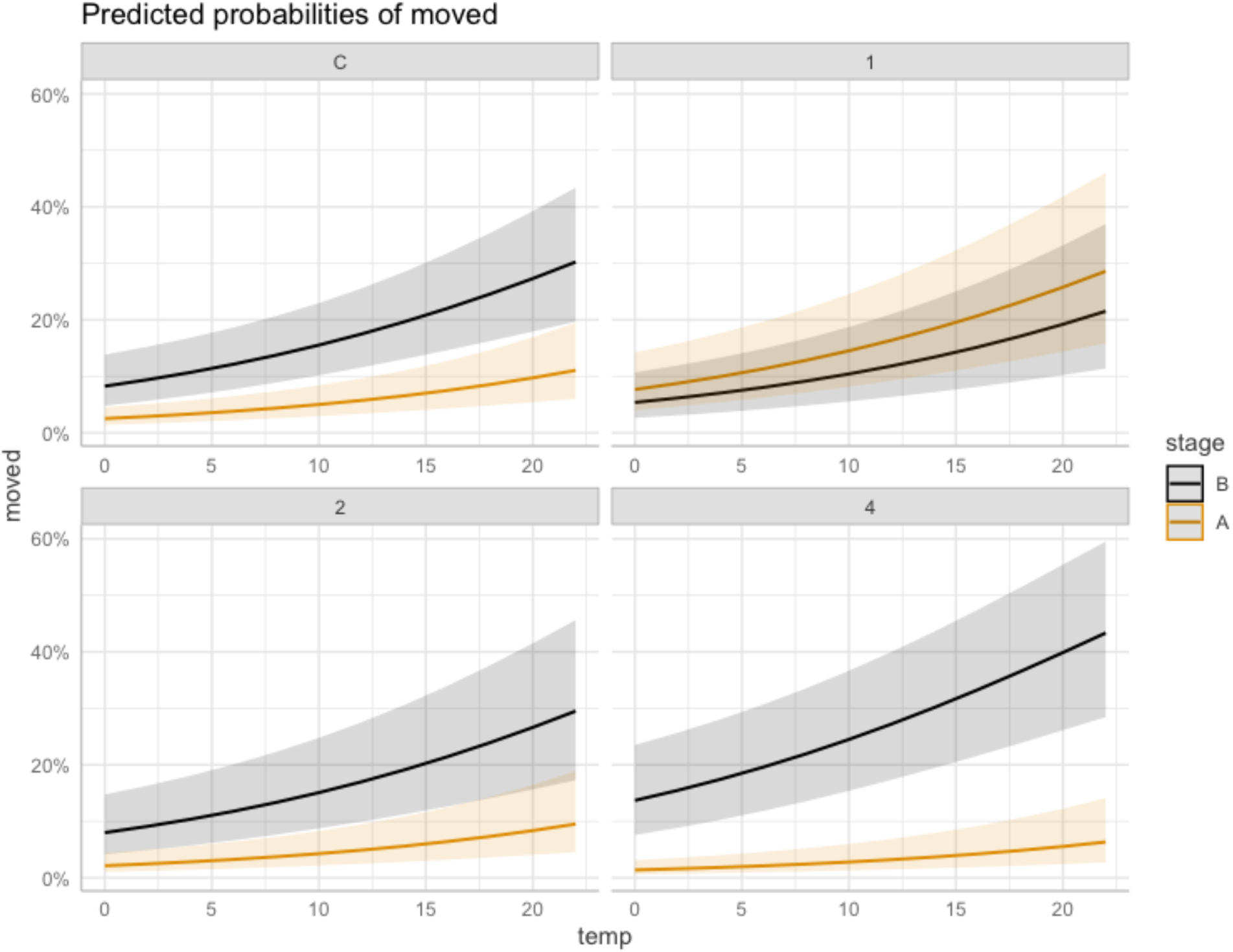
Predicted probabilities of moving compared to not moving in the Before (B) and After (A) stages of the BACI for the Control (C) individuals and those exposed to 1, 2, or 4 bricks.

Disturbed snails appear to show two distinct responses to disturbance. Four disturbed snails did not move at all “After”. The disturbed snails changed their behaviours dramatically compared to the control snails and themselves “Before”, with most avoiding the corner with the brick (Fig. 2, Appendix 1). No snails showed selection or avoidance of the enclosure edge “Before” (Fig. 2, Appendix 1), but “After” those exposed to one or two bricks increasingly selected for the edge and those exposed to four bricks strongly avoided the edge (Fig. 2, Appendix 1).

**Figure 2.**
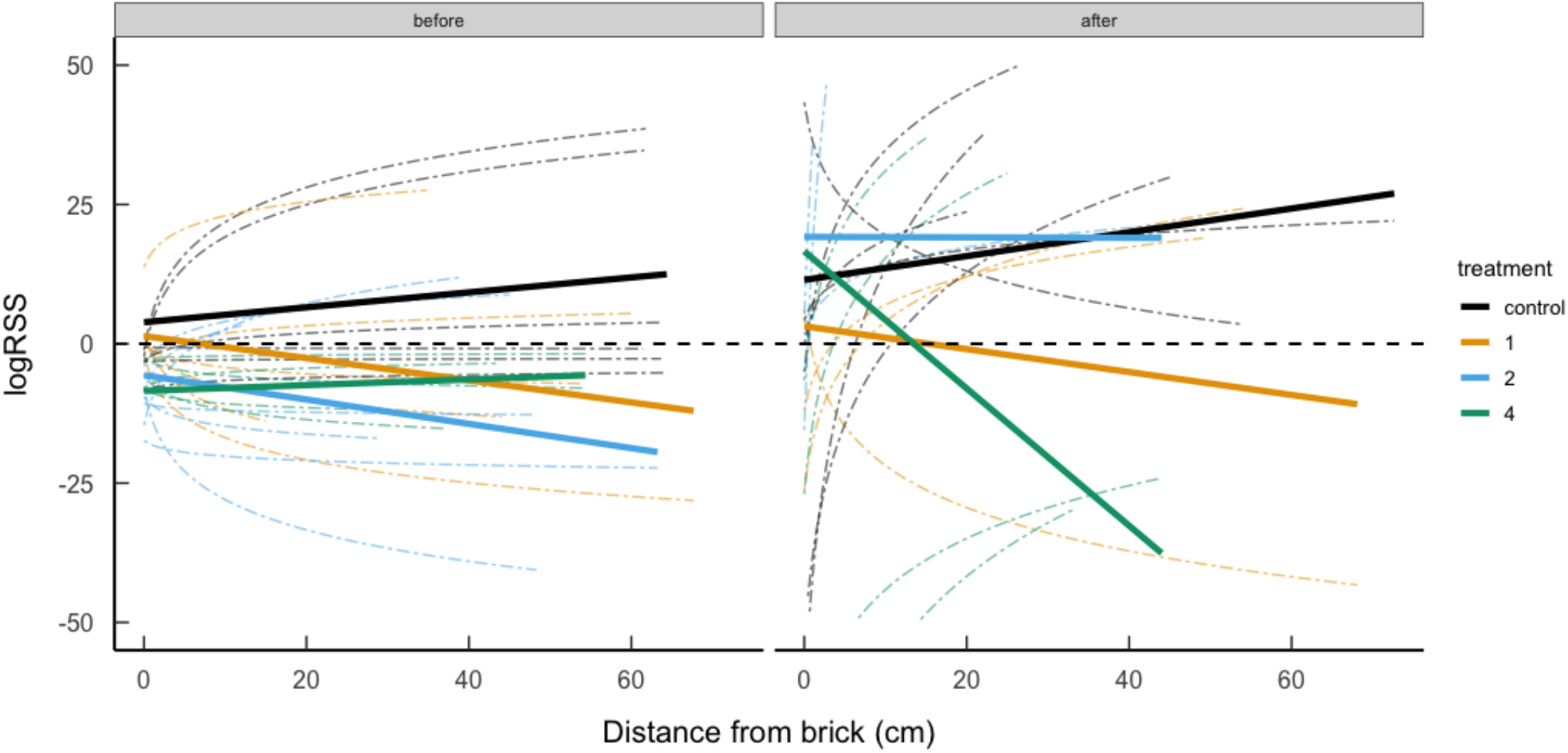
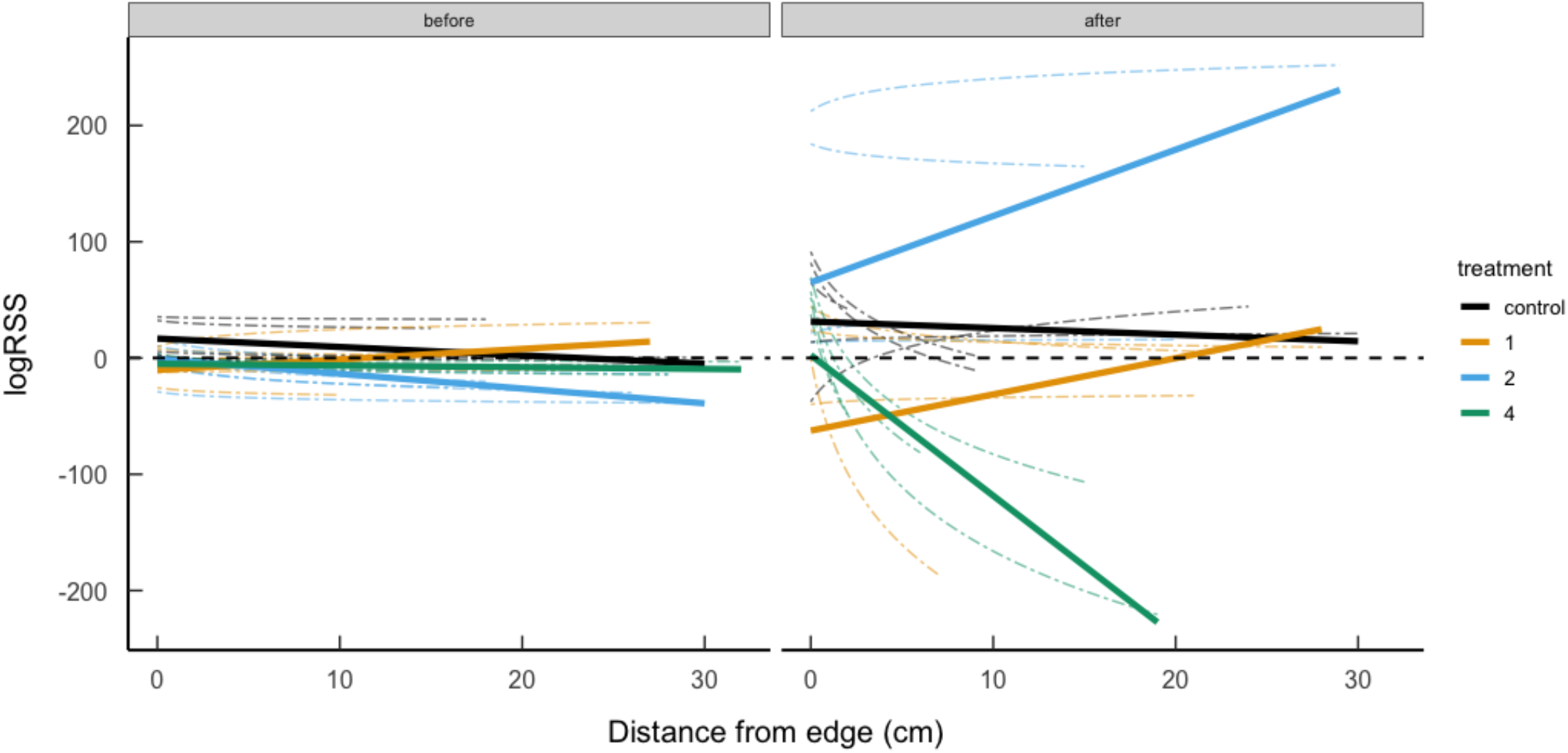
The relative selection strength (RSS) for increasing distances from the corner that would contain a brick (top panel) or the enclosure edge (bottom panel) “After” a disturbed treatment for snails “Before” disturbance and in the undisturbed and disturbed treatments after a brick was added. Dashed lines represent individuals, and solid lines represent group means.

Similarly, no snails adjusted their speed compared to where the brick would be “Before”, though there was some variation in speed relative to the enclosure edge (Fig. 3, Appendix 2). However, all disturbed treatments showed a lot of change in movement “After”, with those exposed to 4 bricks moving significantly faster the farther they are from the brick and enclosure edge (Fig. 3, Appendix 2)

**Figure 3.**
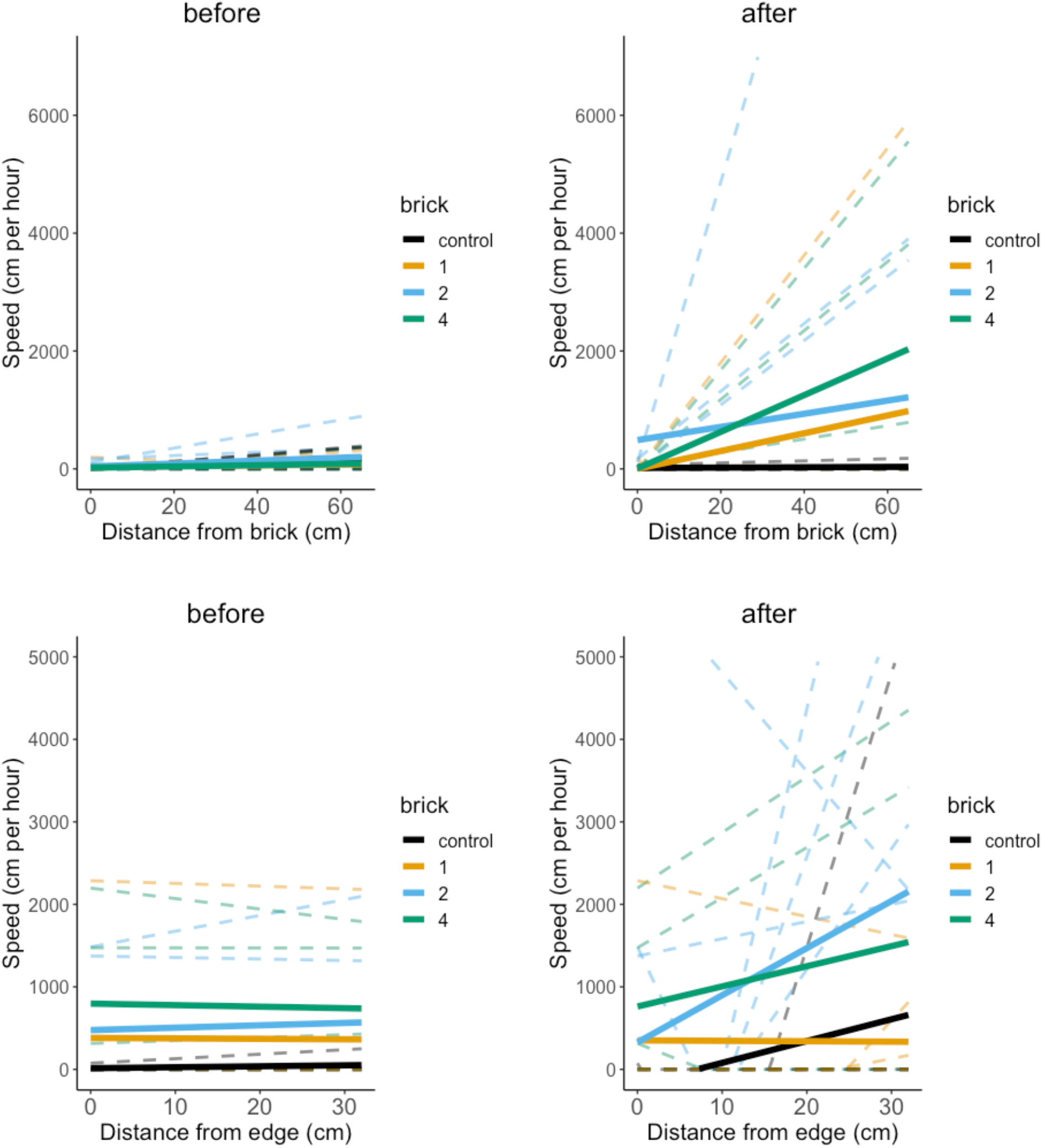
Change in mean speed between the “Before” and “After” stage for snails that were undisturbed and disturbed.

## Discussion

Animals are predicted to alter their movement behaviour in response to risk (Frid & Dill, 2002), though the way they alter this can vary between species. As anthropogenic alterations to habitats such as roads are constructed, habitat loss and fragmentation has become an increasingly prevalent issue that affects the distribution and movement of many animal taxa (Belmeziti et al., 2018; Tucker et al., 2018). We used an iSSA including both habitat and movement parameters to characterize the behavioural responses of grove snails exposed to a physical human disturbance in their habitat. Overall, we found that snails showed a split response to human disturbance; they either stopped moving entirely or moved faster than snails that were not disturbed. Snails, also selected to be farther from the added disturbance. These behavioural changes were greater at higher levels of disturbance. The use of experimental mesocosms permitted us to carefully manipulate edge and disturbance variables while maintaining natural environmental conditions, allowing for models that could disentangle the probable causes behind the behavioural responses observed. Through these models we found evidence that grove snails can perceive edges and respond to them as a risk by altering their movement and space use. Additionally, there was evidence that grove snails can modify their behavioural response as the degree of risk varies, as different behaviours were observed in snails exposed to larger numbers of bricks.

Furthermore, a comprehensive review of species’ movement relative to human disturbance (not including invertebrates) found that the dominant strategy for mammals was to decrease their speed relative to human footprint (Tucker et al., 2018). However, alternate strategies may also compete. For example, our fine-scale approach suggests a bimodal response to disturbance: hunkering down or zooming around. Avoiding risk by reducing movement and seeking cover is commonly observed in animals with low mobility (Denny, 1980). For example, infants of many species are less mobile than conspecific adults, and employ the strategy of hiding to avoid predation (Mabille & Berteaux, 2014). In contrast, highly mobile elk increase movement rates when near a risky area as individuals seek cover from the disturbance (Prokopenko et al., 2017). Although the binary response may be a function of sample size, it highlights the importance of considering alternate adaptive strategies. For example, moving less may minimize encounters with disturbance, but also moving more with a shifted activity budget may also change the manner animals interact with disturbance (Ritzel & Gallo, 2020). Despite our small sample size, we have reasonably large effect sizes, which gives us confidence that our results reflect meaningful strategies employed by snails. Moving faster may not be the dominant response to human disturbance, but having at least some of the population become more mobile may be critical for species’ survival in the Anthropocene.

Anthropogenic disturbance typically accumulates on a landscape (Russell et al., 2021). Responses to increased disturbance may be non-linear, whereby animals can tolerate lower levels of environmental change. However, as effects accumulate, tipping or switch points may occur (Johnson, 2013; Trombulak & Frissell, 2000). Aware of these phenomena, we designed our experiment to simulate accumulating disturbance. Indeed, snails exhibited little response in movement or selection to our lowest two levels of disturbance. In contrast, however, the highest level of disturbance elicited strongest avoidance and faster movement. Grove snails may attenuate their response to disturbance because they have limited mobility and movement is energetically costly (Denny, 1980; Nicolai & Ansart, 2017). This difference in response to varying degrees of risk has also been described in other gastropod species, suggesting a trade-off between risk avoidance and energy cost (Dalesman et al., 2006).

One promise of movement ecology is the likelihood that its inference derives from mechanism (Nathan et al., 2008). But the majority of movement ecology has been enabled by technology and continues to occur in systems that are difficult to manipulate -- manipulations being central to accessing a mechanistic understanding (Holyoak et al., 2008). We capitalize on the snail system to perform replicated treatments within a movement ecology framework. Our experimental setup demonstrated that disturbing natural habitat changes the behavior of individuals. The experiment coupled with models that integrate movement and habitat selection highlighted the importance of individual variation in response to disturbance. Within the individual responses, a bimodal strategy emerged for snails of either ceasing movement or moving significantly more in response to disturbance. Analogous to our expectations from more complex systems, we also observed a ‘tipping point’ or threshold where disturbance appears to have a marked behavioral effect. We hope that our example and our findings foster further experimental movement ecology, to support and complement ongoing observational inferences.

## Acknowledgements

We respectfully acknowledge the territory in which data were collected and analyzed as the ancestral homelands of the Beothuk and the Island of Newfoundland as the ancestral homelands of the Mi’kmaq and Beothuk. We thank all members of the Wildlife Evolutionary Ecology Lab, including K. Kingdon, J. Balluffi-Fry, I. Richmond, J. Hogg, J. Kennah, J. Aubin, J. Hendrix, Q. Webber, S. Boyle, and L. Newediuk for their reviews. We especially thank Mr. Denief for letting us take over his backyard for several months. Funding for this study was provided by a NSERC Discovery Grant to EVW.

## Data Availability

Data and code will be made available upon publication.

## Appendix 1

Model output of the fixed effects from the iSSA examining habitat selection “Before” and “After” added brick disturbance.

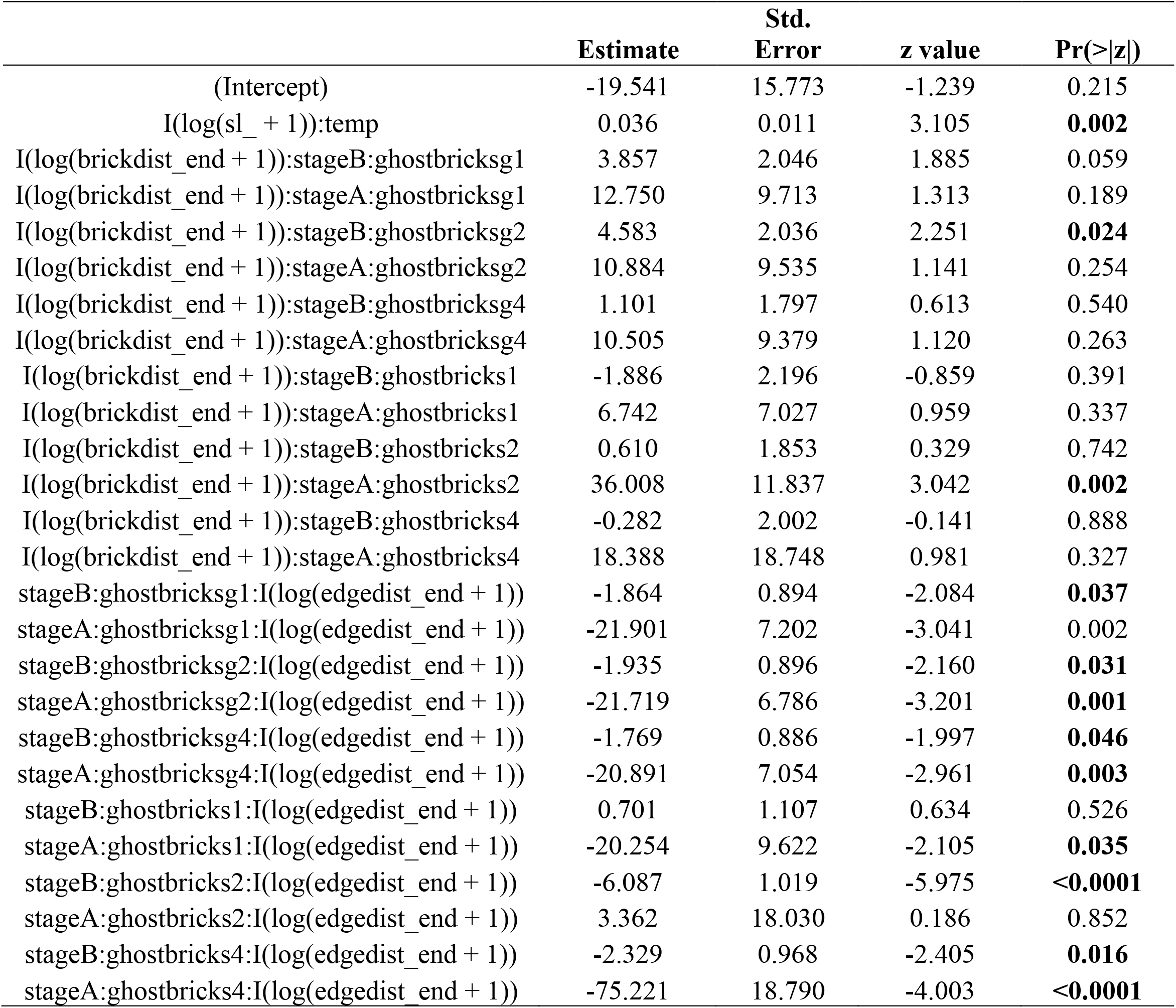

## Appendix 2

Model output of the fixed effects from the iSSA examining movement “Before” and “After” added brick disturbance.

|  | Estimate | Std. Error | z value | Pr(>\|z\|) |
| --- | --- | --- | --- | --- |
| (Intercept) | -2.266 | 14.163 | -0.160 | 0.873 |
| I(log(sl_ + 1)):temp | 0.449 | 0.025 | 17.823 | **<0.0001** |
| I(log(sl_ + 1)):tod_start_night | 6.362 | 0.242 | 26.262 | **<0.0001** |
| I(log(sl_ + 1)):stageB:ghostbricksg1 | -9.035 | 4.769 | -1.895 | 0.058 |
| I(log(sl_ + 1)):stageA:ghostbricksg1 | -16.572 | 6.119 | -2.708 | **0.007** |
| I(log(sl_ + 1)):stageB:ghostbricksg2 | -7.712 | 4.444 | -1.736 | 0.083 |
| I(log(sl_ + 1)):stageA:ghostbricksg2 | -12.736 | 5.200 | -2.449 | **0.014** |
| I(log(sl_ + 1)):stageB:ghostbricksg4 | -6.561 | 4.017 | -1.633 | 0.102 |
| I(log(sl_ + 1)):stageA:ghostbricksg4 | -4.911 | 4.228 | -1.162 | 0.245 |
| I(log(sl_ + 1)):stageB:ghostbricks1 | -5.048 | 4.593 | -1.099 | 0.272 |
| I(log(sl_ + 1)):stageA:ghostbricks1 | -12.799 | 4.759 | -2.690 | **0.007** |
| I(log(sl_ + 1)):stageB:ghostbricks2 | -11.230 | 4.166 | -2.695 | **0.007** |
| I(log(sl_ + 1)):stageA:ghostbricks2 | -5.589 | 4.891 | -1.143 | 0.253 |
| I(log(sl_ + 1)):stageB:ghostbricks4 | -7.818 | 5.437 | -1.438 | 0.150 |
| I(log(sl_ + 1)):stageA:ghostbricks4 | -17.068 | 8.773 | -1.946 | 0.052 |
| I(log(sl_ + 1)):stageB:ghostbricksg1:I(log(brickdist_start + 1)) | -0.155 | 1.469 | -0.106 | 0.916 |
| I(log(sl_ + 1)):stageA:ghostbricksg1:I(log(brickdist_start + 1)) | -14.634 | 38.387 | -0.381 | 0.703 |
| I(log(sl_ + 1)):stageB:ghostbricksg2:I(log(brickdist_start + 1)) | -0.613 | 1.415 | -0.433 | 0.665 |
| I(log(sl_ + 1)):stageA:ghostbricksg2:I(log(brickdist_start + 1)) | 8.958 | 38.481 | 0.233 | 0.816 |
| I(log(sl_ + 1)):stageB:ghostbricksg4:I(log(brickdist_start + 1)) | -1.057 | 1.313 | -0.805 | 0.421 |
| I(log(sl_ + 1)):stageA:ghostbricksg4:I(log(brickdist_start + 1)) | 1.264 | 38.510 | 0.033 | 0.974 |
| I(log(sl_ + 1)):stageB:ghostbricks1:I(log(brickdist_start + 1)) | -2.596 | 1.529 | -1.697 | 0.090 |
| I(log(sl_ + 1)):stageA:ghostbricks1:I(log(brickdist_start + 1)) | 15.574 | 42.193 | 0.369 | 0.712 |
| I(log(sl_ + 1)):stageB:ghostbricks2:I(log(brickdist_start + 1)) | 1.422 | 1.533 | 0.928 | 0.354 |
| I(log(sl_ + 1)):stageA:ghostbricks2:I(log(brickdist_start + 1)) | 58.966 | 41.475 | 1.422 | 0.155 |
| I(log(sl_ + 1)):stageB:ghostbricks4:I(log(brickdist_start + 1)) | 0.872 | 1.942 | 0.449 | 0.653 |
| I(log(sl_ + 1)):stageA:ghostbricks4:I(log(brickdist_start + 1)) | -45.253 | 56.696 | -0.798 | 0.425 |
| I(log(sl_ + 1)):stageB:ghostbricksg1:I(log( edgedist_start + 1)) | 0.818 | 1.031 | 0.794 | 0.427 |
| I(log(sl_ + 1)):stageA:ghostbricksg1:I(log( edgedist_start + 1)) | 54.535 | 87.097 | 0.626 | 0.531 |
| I(log(sl_ + 1)):stageB:ghostbricksg2:I(log( edgedist_start + 1)) | 0.842 | 1.074 | 0.784 | 0.433 |
| I(log(sl_ + 1)):stageA:ghostbricksg2:I(log( edgedist_start + 1)) | 41.267 | 85.880 | 0.481 | 0.631 |
| I(log(sl_ + 1)):stageB:ghostbricksg4:I(log( edgedist_start + 1)) | 0.405 | 1.049 | 0.386 | 0.700 |
| I(log(sl_ + 1)):stageA:ghostbricksg4:I(log( edgedist_start + 1)) | 33.228 | 84.945 | 0.391 | 0.696 |
| I(log(sl_ + 1)):stageB:ghostbricks1:I(log( edgedist_start + 1)) | 2.372 | 1.346 | 1.762 | 0.078 |
| I(log(sl_ + 1)):stageA:ghostbricks1:I(log( edgedist_start + 1)) | 25.934 | 81.647 | 0.318 | 0.751 |
| I(log(sl_ + 1)):stageB:ghostbricks2:I(log(edgedist_start + 1)) | 0.829 | 1.093 | 0.759 | 0.448 |
| I(log(sl_ + 1)):stageA:ghostbricks2:I(log(edgedist_start + 1)) | -138.561 | 86.974 | -1.593 | 0.111 |
| I(log(sl_ + 1)):stageB:ghostbricks4:I(log(edgedist_start + 1)) | -0.965 | 1.561 | -0.618 | 0.536 |
| I(log(sl_ + 1)):stageA:ghostbricks4:I(log(edgedist_start + 1)) | -25.386 | 98.832 | -0.257 | 0.797 |

